# Barbary ground squirrels do not have a sentinel system but instead synchronise vigilance

**DOI:** 10.1101/2020.04.24.055707

**Authors:** Annemarie van der Marel, Jane M. Waterman, Marta López-Darias

## Abstract

Coordinated behavior, such as hunting in lions and coordinated vigilance as antipredator behavior, are examples of benefits of group-living. Instead of asynchronous vigilance, some social species synchronize their vigilance bouts or take turns acting as sentinels. To increase our knowledge on the evolution of vigilance behavior, we studied whether vigilance is coordinated in Barbary ground squirrels, *Atlantoxerus getulus*. We show that vigilance was synchronized instead of taking turns. Multiple non-mutually exclusive hypotheses could explain synchronization: Barbary ground squirrels may perch because 1) neighbors are perched (copying effect), 2) perch synchrony may be an emergent property of the ecology as all squirrels may be satiated at the same time (collective behavior), or 3) the benefits are large in terms of evading ambush predators and scanning effectiveness (watch each other’s back). Particularly, in habitats where the field of view is obstructed by man-made structures and multiple individuals may be necessary to watch for terrestrial predators, synchronized vigilance may have greater fitness benefits than sentinel behavior. We conclude that it is essential to test assumptions of coordination and, thus, to analyze coordination to describe sentinel systems.

**Significance Statement:** Vigilance behavior can be vital to an animal’s survival. Taking turns acting as sentinels or synchronizing vigilance bouts reduces the cost of the trade-off between feeding and predation risk. A sentinel system assumes that sentinels are vigilant from raised positions, warn group members of danger, and alternate vigilance bouts. However, the assumption of alternating vigilance bouts remains poorly tested. We tested this assumption in invasive Barbary ground squirrels. We found that instead of alternating, individuals synchronized their vigilance bouts. Perch synchrony may be 1) a response to perching group members (copying effect), 2) an emergent property of the species’ ecology, and 3) an adaptation to anthropogenically altered habitats (watch each other’s back).

## Introduction

Vigilance is essential for diurnal species to avoid predation resulting in a trade-off with foraging (Bednekoff and Lima 1998). Individuals are expected to balance the trade-off to increase their survival chances. In diurnal prey species, the cost of vigilance (less time available for foraging) can be reduced by performing nonexclusive or low-quality vigilance, i.e., individuals are vigilant while performing another behavior (e.g. Cape ground squirrels, *Xerus inauris*, (Unck et al. 2009), or eastern grey kangaroos, *Macropus giganteus*, (Favreau et al. 2015)). Low-quality vigilance is often seen in habitats where the risk imposed by predators is relatively low (Unck et al. 2009; Périquet et al. 2012). In riskier habitats, individuals increase their time spent in exclusive or high-quality vigilance –vigilance without performing any other behavior– while reducing the time spent foraging (Unck et al. 2009; Périquet et al. 2012). Thus, multi-tasking is one way to reduce the cost.

Another way to mitigate the foraging/vigilance trade-off cost is by taking turns acting as sentinels (nonoverlapping vigilance bouts) or by synchronizing vigilance (overlapping vigilance bouts) among group members (Beauchamp 2015). Synchronization is frequent both in birds and mammals (Ebensperger et al. 2006; Pays et al. 2007a, b, 2009, 2012; Öst and Tierala 2011; Podgórski et al. 2016; McDougall and Ruckstuhl 2018). Synchronization may occur because individuals are vigilant at the same time due to the same external stimulus (e.g., a predator, termed induced vigilance, Blanchard and Fritz 2007) or because individuals copy group members’ behavior (allelomimetic copying effect, Deneubourg and Goss 2010).

Some highly social bird and mammal species living or foraging in open arid habitats perform sentinel behavior (Rasa 1977; Ferguson 1987; McGowan and Woolfenden 1989; Clutton-Brock 1999; Manser 1999; Wright et al. 2001a; Newbold et al. 2008; Arbon et al. 2020). A sentinel system assumes that sentinels warn group members of danger (Bednekoff 1997) and that there is a continuous turn-over of who is sentinel, but there is no change in sentinel number (McGowan and Woolfenden 1989; Bednekoff 2015). In species with a sentinel system, one or two group members are always vigilant because individuals forego feeding and take turns performing high-quality vigilance from raised positions, which allows for an efficient way to share information about predators in a foraging group (Rasa 1987; Bednekoff 1997; Beauchamp 2015). Sentinels appear to be more effective at detecting predators due to their more focused vigilance and their elevated positions (Ridley et al. 2013), which may also allow them to retreat to cover before group members. Plus, the decision to become sentinel can be state-dependent; individuals in good body condition can afford to act as sentinels more often than individuals in poor body condition (Wright et al. 2001a, b). Nevertheless, sentinel behavior can be costly as sentinels are more likely to attract the attention of predators, e.g., pied babbler, *Turdoides bicolor*, sentinels incur higher predation risk than non-sentinels (Ridley et al. 2013). Thus, coordination of guarding bouts may be a necessity to divide the costs of being on guard over group members. Since non-sentinels can be located at raised positions or can alarm signal, Bednekoff (2015) argued that the main criterion of a sentinel system is the alternation of the high-quality vigilance bouts, however, coordination was tested in only three species (meerkats, *Suricata suricatta*, superb fairy-wrens, *Malurus cyaneus*, and Florida scrub-jays, *Aphelocoma coerulescens*) and suggested for 14 out of 40 species.

In this contribution we explored vigilance behavior of the Barbary ground squirrels, *Atlantoxerus getulus*, and tested whether it can be considered a sentinel system. *A. getulus* is an appropriate study system as both males and females perform high-quality vigilance (actively scan the environment without performing any other behavior for approximately 40% of their daily activity budget) (van der Marel et al. 2019). Barbary ground squirrels perform high-quality vigilance behaviors either on the ground or away from foraging areas in raised positions (> 30 cm from the ground) (AvdM, pers. obs.). We termed this latter vigilance type ‘perch behavior’. Perch behavior seems similar to sentinel behavior as perched individuals are located at raised positions and alarm call to inform group members of danger (van der Marel et al. 2019). Thus, although Barbary ground squirrels meet two criteria of a sentinel system (van der Marel 2019; van der Marel et al. 2019), such behavior cannot be considered as sentinel behavior unless it is coordinated by individuals taking turns (following Bednekoff 2015).

Our objectives were 1) to describe perch behavior in this species, 2) to investigate whether perch behavior was costly, and 3) if Barbary ground squirrels take turns acting as sentinels. First, we explored details of perch and foraging behavior to better understand the foraging/vigilance trade-off in this species. We also tested whether perch behavior was induced by predators. Second, we hypothesized that perch behavior would be costly because squirrels have less time to forage and attract the attention of predators when perched. Third, we hypothesized that Barbary ground squirrels coordinated perch behavior by taking turns acting as sentinels, which we explored as differences in proportions (p_obs_-p_exp_) following Beauchamp (2015), and tested whether coordination was dependent on predator presence.

## Materials and methods

### Study sites, species, and trapping protocol

We studied an invasive population of Barbary ground squirrels in the northwest of Fuerteventura, Canary Islands, Spain (28°34′60” N, 13°58′0” W), from March through July 2014, January through July 2015, and January through June 2016. Fuerteventura has an arid climate and semi-desert habitats characterized by xerophytic scrubland and ravines caused by erosion (del Arco Aguilar et al. 2010). Dams and rock walls, built to make terraces for land cultivation and to fence properties, characterize island landscapes and also field sites, and function as shelters and perches for the Barbary ground squirrels to watch for predators (López-Darias and Lobo 2008; van der Marel et al. 2019). The study area consisted of three study sites on average 0.73 km apart from one another (Site 1: 283558.1N, 135949.15W; Site 2: 283536.81N, 135948.71W; and Site 3: 283538.51N, 140001.41W). The sites differed in the number of squirrels and size (density of squirrels per hectare was 14.15, 9.19 and 7.24 for sites 1, 2 and 3, respectively, van der Marel et al. 2019, 2020), but did not represent distinct squirrel populations (van der Marel et al. 2020). Over the three years, we sighted a total of 175 aerial and 96 terrestrial predators and the number of predator sightings was similar across all three sites (van der Marel et al. 2019). The main predators of Barbary ground squirrels on Fuerteventura are the native Eurasian buzzard, *Buteo buteo* (Gangoso et al. 2006), the invasive feral cat, *Felis catus* -- the only terrestrial predator-- (Medina et al. 2008), and occasionally the common raven, *Corvus corax* (Gangoso et al. 2006) and the common kestrel, *Falco tinnunculus* (López-Darias and Lobo 2008). We observed five successful (out of 10) aerial predator attacks and one successful (out of 4) terrestrial predator attacks (van der Marel et al. 2019). As the sites were close to one another and only differed in squirrel density, we decided not to include site as a random factor in any of our analyses.

As part of a broader research project, we continuously monitored the population of the study area by trapping Barbary ground squirrels weekly throughout the three field seasons (procedures are described in Piquet et al. 2018; van der Marel et al. 2019, 2020, 2021). For individual identification, we PIT-tagged and dorsal dye marked individuals. We additionally measured squirrel body mass and hindfoot length (mm) as proxy measurements for body size (Schulte-Hostedde et al. 2001), using a spring scale (± 5 g; Pesola AG, Baar, Switzerland) and a Digimatic Plastic Caliper (Mitutoyo Corporation, Kawasaki, Japan), respectively. We distinguished adult and subadult males and females by the state of their primary sexual characters (scrotum or vulva and nipples, (van der Marel et al. 2020, 2021a)). Subadults are over six months old and have reached adult body size, but males do not have descended testes and females do not have swollen vulva and nipples during the mating season (i.e., days when females are in estrus). The mating season is distinct as males regress their scrotum once all females have mated (van der Marel et al. 2021a). Males have been observed to travel between sites only during the mating season; otherwise, adult males and females of all ages stay within a site (AvdM, pers. obs.).

Barbary ground squirrels are social because individuals share sleeping burrows and show spatiotemporal overlap and cohesiveness (van der Marel et al. 2020). Females are the philopatric sex, i.e. they share sleeping burrows with related adult females (1-8 females), whereas males disperse before sexual maturation and share sleeping burrows with unrelated males and subadults of either sex (1-16 individuals) (van der Marel et al. 2020). Female sleeping burrow group composition is more stable than for males; but throughout the day, males and females of different sleeping burrow associations can be active in the same area in a site, showing fission-fusion dynamics (van der Marel et al. 2020). Therefore to accommodate for the fluid group compositions, we used two group definitions: 1) individuals sharing a sleeping burrow together (nighttime social groups) and 2) a set of individuals that were in each other’s visible range during an observation period (daytime social group) (Stankowich 2003; van der Marel et al. 2019). We did not consider auditory range as part of our group definition as we heard alarm calls from different sites and we do not know at what distance squirrels can hear alarm calls. In our study area, squirrels prefer small ravines with old lava tunnels that function as burrows or areas with rock walls that formed terraces for land cultivation (López-Darias and Lobo 2008; van der Marel et al. 2020).

### Behavioral observations

To characterize the perch behavior of Barbary ground squirrels, we used 10 min scan samples throughout each behavioral observation period (Altmann 1974). It was not possible to record data blind because our study involved focal animals in the field. We conducted behavioral observations during all three field seasons between 9:30 h GMT and 2 h before sunset, i.e., when the squirrels are active above ground (Machado 1979; van der Marel et al. 2019). We observed the squirrels within each site from elevated areas and roads, at a distance of approximately 50 m, which did not affect the squirrels’ behavior. The human observers that visited the sites were not perceived as a threat to the squirrels as the squirrels did not change their behavior when the observers arrived (AvdM, pers. obs). Primarily, we recorded perch behavior, low-quality vigilance (vigilance at any location that lasted < 30 s or was performed while doing other behaviors), and feeding behaviors (actively searching for, manipulating, and/or ingesting a food item). We also recorded whether any predators or other external stimuli were observed in a scan, as animals may respond to the same external stimulus (Beauchamp et al. 2012; McDougall and Ruckstuhl 2018). To enter behavioral data, we used the spreadsheet program Numbers (Apple, Cupertino, CA, USA) on an iPod (Apple, Cupertino, CA, USA) from 2014 until June 2015, and thereafter Prim8 Software (McDonald and Johnson 2014) on an Android phone (Motorola Droid A850).

### Statistical analyses

For all our analyses, we selected individuals that were observed for at least 50 min over a minimum of five observation periods (Edwards and Waterman 2011). We excluded days that females were in estrus (the mating season) because male behavior changed as they search competitively for estrous females, similar to Cape ground squirrels (Waterman 1998). We performed all analyses in R version 3.4.1 (R Core Team 2017) and we made the figures using “ggplot2” package version 3.0.0 (Wickham 2016). We set statistical significance to the level of 0.05. Unless otherwise indicated, we denoted averages as mean ± standard error (SE).

#### General description of perch behavior

To better understand the trade-off between foraging and perch behavior (costs of perch behavior), we used scan data recorded across all three field seasons to calculate the distance to foraging areas, the distance from cover while foraging, the number of squirrels foraging per scan, and the distance between foraging and perching group members. We calculated the distance from sleeping burrows to foraging areas and from foraging areas to cover (rock walls/burrows) using Google Earth. We calculated the distance between group members per scan for nighttime social group members and for daytime social group members as individuals from the same nighttime social group may stay closer together during the day. For this analysis, we excluded individuals that were out of sight and 33 additional observations with distances over 200 m as these were at the extremes at which we could confidently identify the squirrels. To investigate whether all individuals performed perch behavior, we calculated the percentage of individuals that perched from all marked individuals during the three field seasons. We analyzed whether the ground squirrels increased their perch behavior when predators were observed *vs*. not observed using a Mann-Whitney *U* test. If we saw an aerial predator in a scan, that complete scan was considered to have a predator. If we saw a terrestrial predator, we considered the complete observation session to have a predator, as cats were the ambush predators in our study sites.

#### Costs of perch behavior

We tested if any costs were associated with perch behavior using the full dataset (2014-2016). As dependent variable we used the proportion time spent perching, which followed a beta distribution. As independent variables we used the proportion of time spent feeding, body condition and age at which the squirrels disappeared (survival age). We measured body condition by taking the residuals from a linear regression of log body mass (g) on log hind foot length (mm) (Labocha et al. 2014; Piquet et al. 2018). We calculated survival age as the age the squirrels lived until they disappeared (which entails death as the adult squirrels do not disperse from their site). Survival age may be an underestimate as we did not have the date of birth or the year the squirrels disappeared of all squirrels because our field season was only 3 years in duration. We tested for multicollinearity of our variables using the variance inflation factor (vif) function in the “car” package (Fox and Weisberg 2011). None of our variables showed a vif value above 3, so we included all variables. We used ID and year as random factors. We used glmmADMB (Bolker et al. 2012; Fournier et al. 2012) and we performed a likelihood ratio test (LRT) between the model and a null model using the ‘lmtest’ package version 0.9-36 (Zeileis 2002).

#### Perch behavior coordination analyses

In our coordination analyses, we only used scan data recorded with the Prim8 software, as this software measured the exact duration of perch behavior instead of the number of times the behavior was observed. So, we excluded instances where perch events were shorter than 30 s. To measure coordination of perch bouts, we compared the observed proportion of time when at least one individual was perched per group with the expected proportion under the assumption of independent vigilance following the descriptions in, e.g., Pays et al. (2007a), Öst and Tierala (2011) or Beauchamp (2015), which is the opposite from what is described in Bednekoff (2015). If the difference in proportions (p_obs_-p_exp_) did not differ from 0, then individual perch bouts were independent; if the difference in proportions was greater than 0, then perch bouts were sentinel; and if the difference was smaller than 0, then perch bouts were synchronized (Beauchamp 2015). We measured the observed time as the total time spent perched divided by the total time observed for each individual in the group. We measured the expected time as 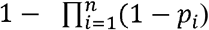, where *n* is the sample size and *p_i_* is the proportion of time that individual *i* is perched, i.e., total time spent perched divided by the total time observed for that individual (Pays et al. 2007a; Öst and Tierala 2011; Beauchamp 2015). We calculated the duration of observed and expected time when at least one individual was perched for all individuals, only adult individuals, and only adult females, excluding unidentified and unknown squirrels. We used two different sample size definitions; first, we used the number of observation periods as our sample size (daytime social group), assuming that the individuals seen in an observation period belonged to the same group (see methods section *Behavioral observations*). We used a non-parametric paired test (Wilcoxon signed-rank test with continuity correction) to test for differences between the expected and observed values per observation period. Second, to prevent pseudoreplication, we used the number of nighttime social groups, which we defined as individuals sharing the same sleeping burrow (i.e., sleeping burrow associations), as our sample size. We tested for differences between the expected and observed values using a glmmADMB (Bolker et al. 2012; Fournier et al. 2012) and between the model and the null model using the likelihood ratio test (LRT) from the ‘lmtest’ version 0.9-36 (Zeileis 2002). We used the proportion of time spent perching as dependent variable and the proportion type (observed or expected) as independent variable, with year and social group as random factors. Using social groups as our sample size, we also tested for coordination when predators were (n = 48 scans in 20 observation sessions) and were not (n = 977 scans in 176 observation sessions) observed. For this analysis, we only selected social groups that were observed during at least 2 predator encounters each year (n = 9 social groups). We used a glmmADMB with the proportion of time spent perched as dependent variable, an interaction between proportion type and predator presence as independent variable and year and group ID as random factors.

## Results

### General description of perch behavior

We observed the squirrels on average for 9 ± 7.3 (mean ± SD) scans per observation period (n = 359 observation periods). Upon emergence from their sleeping burrows in the morning (9:30-11:00), the ground squirrels were slow to wake-up, but eventually, they moved away from their sleeping burrows to foraging areas, with the last squirrels immerging in their burrows as early as 15:30 when it was cool and rainy and as late as 20:00 in the summer months (AvdM, pers. obs.). We observed 418 feeding and 1635 perch bouts in the morning (between 9:30 and 12:00), 366 and 1084 during midday (12:00-15:00) and 994 and 1975, respectively, in the afternoon (15:00 until immergence into the sleeping burrows, between 15:30 and 20:00). The proportion of individuals feeding and perching in a 10-minute scan was 0.13 ± 0.22 (mean ± SD, n = 3231 scans) and 0.39 ± 0.34 (mean ± SD, n = 3231 scans), respectively, ranging from no individuals to all individuals performing the behavior. The maximum distance that squirrels moved from sleeping burrows to foraging areas was 185.5 ± 78.2 m (mean ± SD, n = 14) and the average distance to cover when foraging was 3.7 ± 3.1 m (mean ± SD) with a maximum distance of 20 m (n = 154 observations). The distance between individuals within a daytime social group that were observed in the same scan was 41.5 ± 29.7 m (mean ± SD, n = 5450 observations), ranging from 0 to 197 m. Between individuals that belonged to the same sleeping burrow association/nighttime social group, the distance was 36.7 ± 28.0 m (range: 0-187 m, n = 2677 observations). Per scan, the average distance was 22.7 ± 20.9 m (range: 0-139 m, n = 511) between foraging and 44.2 ± 29.7 m (range: 0-197 m, n = 1508) between perching individuals. When selecting for only nighttime social group members, the average distance was 20.7 ± 20.4 m (range: 0-112 m, n = 254) for foraging and 38.8 ± 27.5 m (range: 0-183 m, n = 698) for perching individuals.

Overall, 99.3% of the squirrels (adults, subadults, and juveniles, n = 142) were observed perching. In 32.4% of a total of 3233 scans, just one individual perched, while more than one squirrel was perched in 39.8% of scans. When predators were not observed, one individual perched in 32.9% and more than one squirrel was perched in 38.4% scans of a total of 3023 scans (Fig. 1a). When predators were observed, one squirrel was perched in 24.8% of scans and more than one squirrel in 59.5% of scans (n = 210 scans, Fig. 1a). Individuals perched both in the presence (0.46 ± 0.02, n = 210 scans) and absence of predators (0.38 ± 0.01, n = 3023 scans), although the proportion of individuals perched per scan when a predator was observed was significantly higher than when there was no external stimulus (Mann-Whitney *U* test: *U* = 267477, *P* < 0.001, effect size r = 0.07; Fig. 1b).

**Fig. 1.**
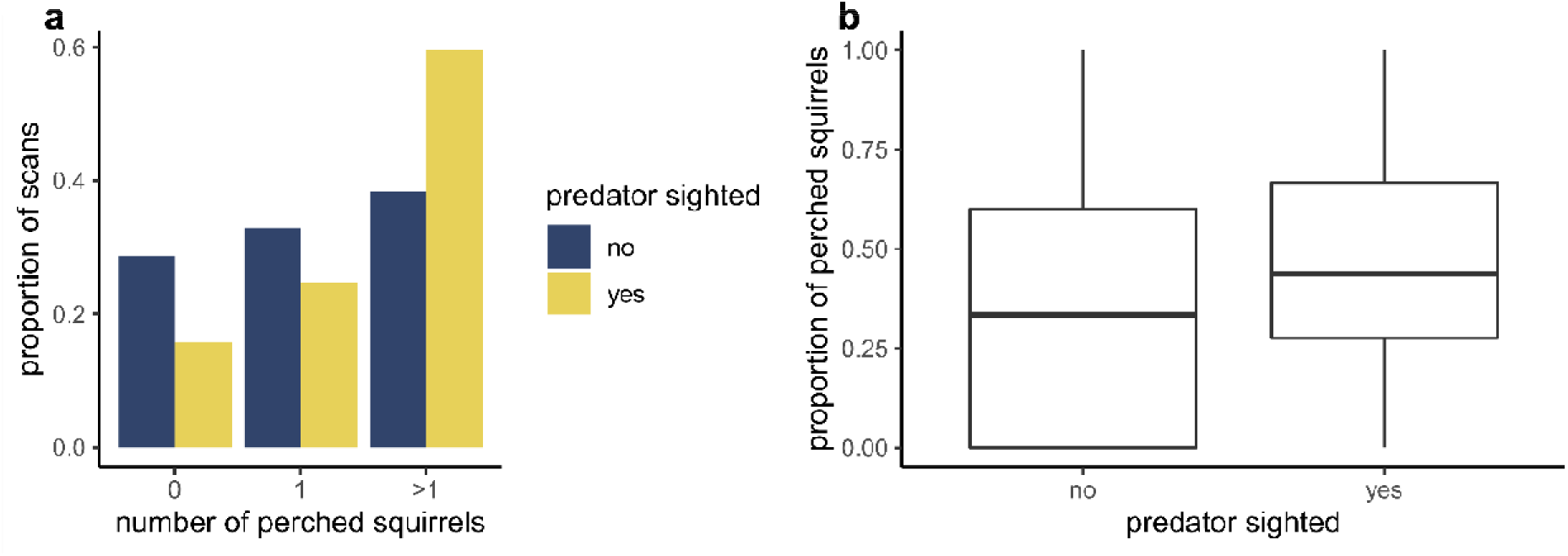
a) The proportion of scans where 0, 1, or more than 1 Barbary ground squirrels were perched when predators were and were not observed (controlled for number of scans), and b) the proportion of perched squirrels when predators were and were not observed with the dark line representing the median, the box edges are the upper and lower quartiles, the whiskers are 50% from the median, and the closed circles are the outliers, calculated as the values smaller or larger than 1.5 times the box length (i.e., upper-lower quantile).

### Costs of perch behavior

We found that costs are associated with the proportion of time spent perching (LRT, Λ = 45.2, *P* < 0.001; Fig. 2). Yet, although we found a negative relation between the amount of time spent perched and foraging (GLMM: −2.38 ± 0.42, p < 0.001, random factors ID = 0.04 and year = 0.05, Fig. 2a), body condition was positively related to time spent perching (GLMM: 0.56 ± 0.19, p = 0.003, Fig. 2b). We did not find a relation between the proportion of time spent perched and survival age (GLMM: −0.03 ± 0.04, p = 0.45, Fig. 2c).

**Fig. 2.**
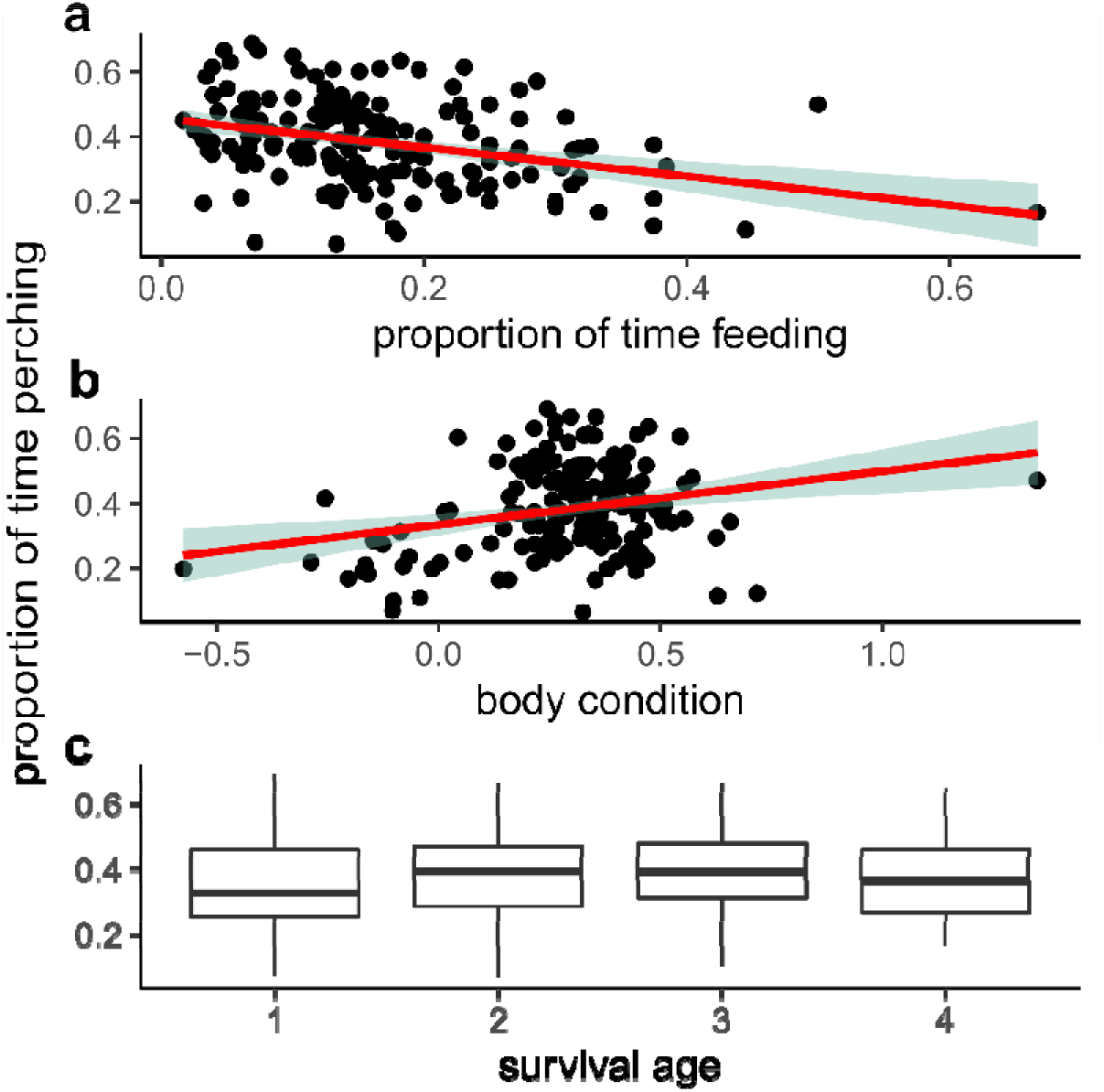
The relationship between the proportion of time perching and a) foraging, b) body condition, and c) survival age. In a and b, the red lines represent the regression line and the shaded areas are the confidence interval. In c, the boxes indicate the inter quartile range (IQR), with the central line depicting the median and the whiskers extending to 1.5*IQR

### Perch behavior coordination analyses

We found that the proportion of observed time that at least one individual in the group (defined as either observation period or sleeping burrow association) was perched (p_obs_) was smaller than the proportion of expected time (p_exp_, Table 1). Using sleeping burrow associations/nighttime social groups, we also found that the observed proportion of time perching was smaller than the expected proportion (GLMM: 1.09 ± 0.23, p < 0.001, random factors year = 0.05 and group = 0.36, Table 1). Using a subset of the data to study the effect of predator presence on proportion of time spent perching, we found that the observed proportion of time perching was also smaller than the expected proportion (LRT, Λ = 28.6, *P* < 0.001; GLMM: 1.66 ± 0.33, p < 0.001, random factors year = 0.11 and group = 0.24). We found no effect of predator presence (GLMM: −0.29 ± 0.35, p < 0.42, Fig. 3), but there was an interaction between the proportion of time spent perched and predator presence, where the difference between the proportions was smaller when predators were present (GLMM: −1.08 ± 0.47, p = 0.02, Fig. 3).

**Table 1.**
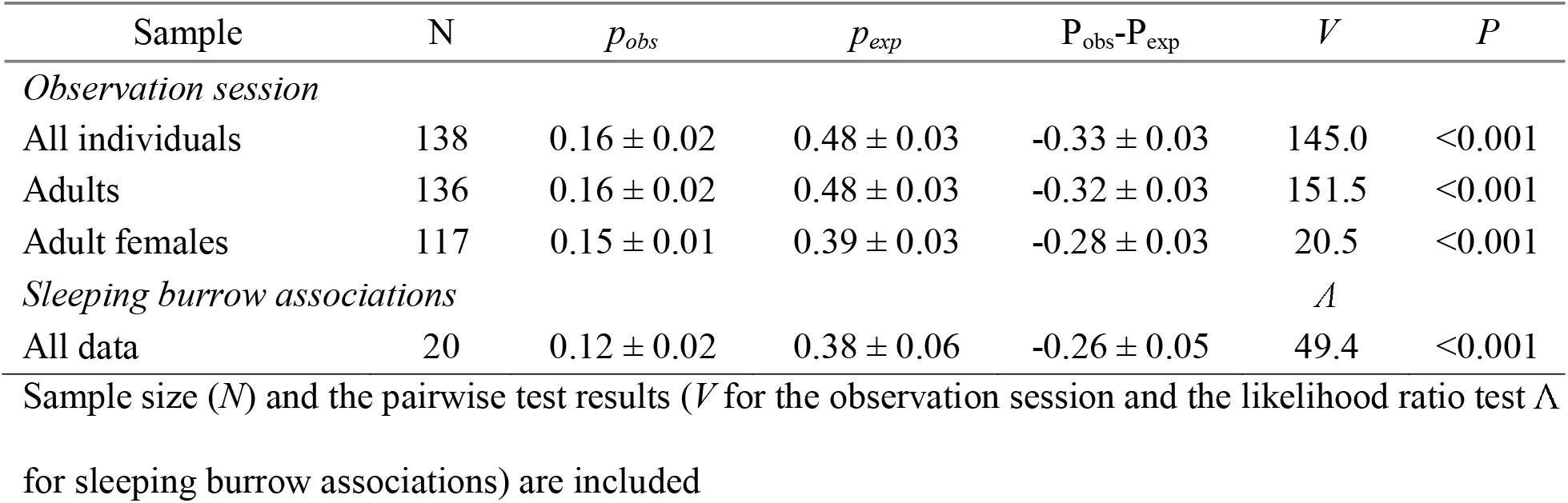
The proportion of observed (p_obs_) and expected (p_exp_) time at least one individual was perched (mean ± SE) per observation period in Barbary ground squirrels, calculated as duration of perch bouts

**Fig. 3.**
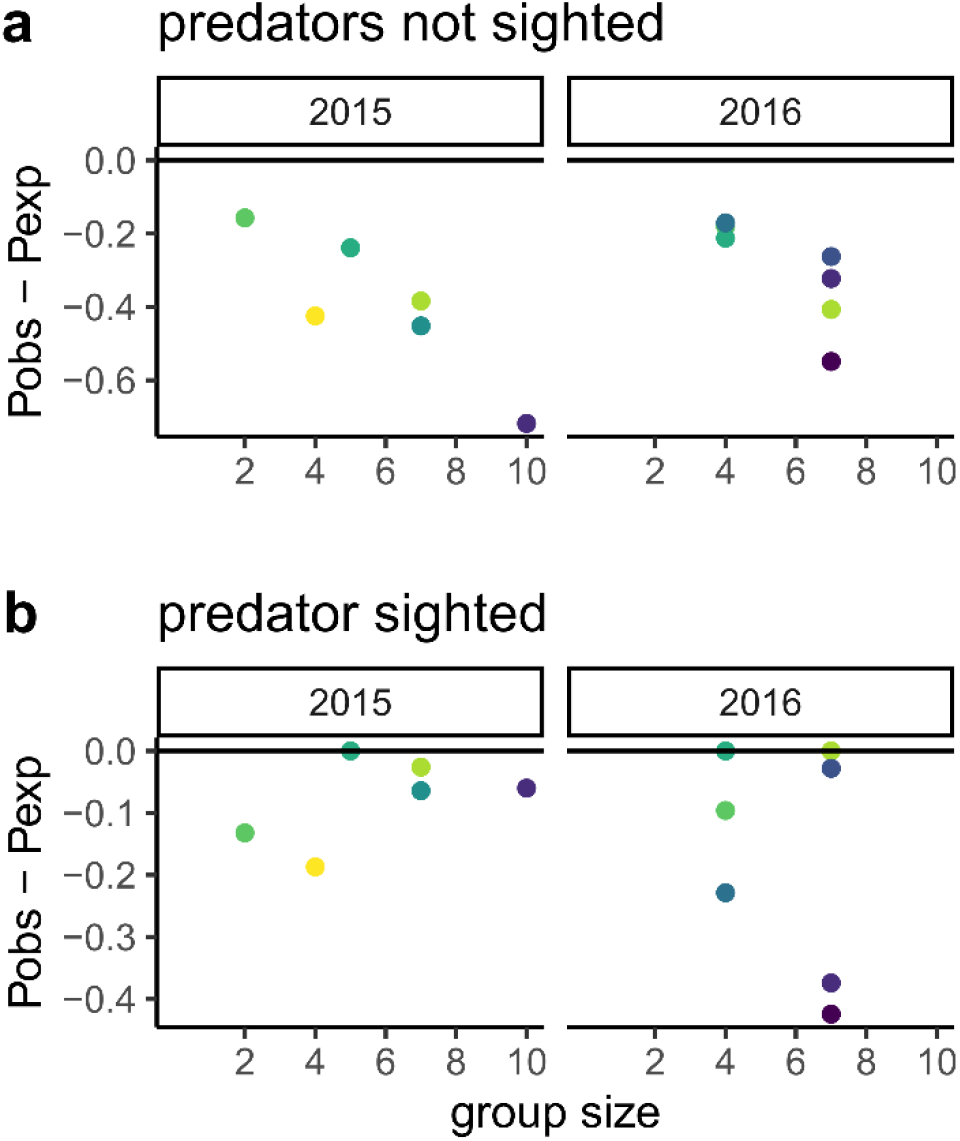
Difference between the observed (p_obs_) and expected (p_exp_) proportions of time when at least one Barbary ground squirrel was perched per group, defined as a sleeping burrow association, when predators a) were not and b) were sighted. Each color represents a different social group (n = 9). When the difference is smaller than 0, vigilance is synchronized

## Discussion

We studied high-quality vigilance (‘perch’ behavior) in Barbary ground squirrels and whether perch behavior was costly and coordinated by taking turns acting as sentinels. Perch behavior was performed in the absence and presence of an external stimulus (a predator), yet perch behavior increased when predators were observed. We found that perch behavior resulted in less time foraging, however this reduction did not result in lower body condition or lower survival chances. When testing the assumption of taking turns, the results may show that coordination of vigilance is absent altogether (Hing et al. 2019), synchronized (e.g., our study; Wang et al. 2020), or switches between synchronization and taking turns dependent on level of disturbance and predation risk (Kong et al. 2021). In Barbary ground squirrels, perch behavior was synchronized because the observed time that at least one individual is perched was smaller than the expected time, resulting in a difference in proportions smaller than zero. Additionally, synchronization occurred when predators were and were not observed.

What are the costs associated with perch behavior? First, we found a negative correlation between proportion of time spent foraging and perching and a positive correlation between the proportion of time spent perching and body condition. We can interpret this using two non-mutually exclusive pathways: the costs of perch behavior are low, as Barbary ground squirrels may have the time and energy to perform perch behavior as an optimal activity, similar to state-dependent sentinel behavior as is seen in suricates, *Suricata suricatta* (Clutton-Brock 1999) or Arabian babblers, *Argya squamiceps* (Wright et al. 2001b). Alternatively, perch behavior is condition dependent as only the individuals in better body condition can afford to perch, which results in an underlying cost to foregoing foraging. Second, although, individuals that were located at raised positions were more likely to attract the attention of predators and may have incurred higher predation risk (Ridley et al. 2013), we did not find that perch behavior was costly due to predators. Barbary ground squirrels performed more perch behavior when predators were present, but the effect size was small and perching did not result in lower survival. The rock walls that are present on Fuerteventura provide not only better lookouts but also more shelter options for the squirrels (van der Marel et al. 2019). Thus, when Barbary ground squirrels are perched at elevated positions, they can attract the attention of predators, but the squirrels are closer to cover than foraging group members and can detect predators earlier. Also, predator diversity is lower in the invasive range (one terrestrial and three aerial predators) compared to the native range (eight terrestrial and two aerial predators, Machado and Domínguez 1982). Despite an underlying cost associated with foregoing foraging, the perching Barbary ground squirrels on Fuerteventura may be able to afford a reduction in foraging behavior due to the available resources or the release of enemies (parasites and predators) on the island compared to the native range (López-Darias 2007; López-Darias and Nogales 2008; Piquet et al. 2018). Thus, this interplay of the cost of perching and the benefit of being closer to cover requires further attention.

Since it seems counterintuitive to be on the lookout at the same time as your group members, the question is: Why perch synchronically? Multiple mechanisms could explain synchronization of perch bouts. First, individuals could simply copy the behavior of neighbors (Pays et al. 2007a). Vigilance is then induced by other vigilant individuals, which may reflect perceived predation risk of the group (Beauchamp 2015). Thus, the squirrels may have a simple rule that when another squirrel is perched, it should perch too, as there might be an ambush predator present. This mechanism is a selfish strategy as it will be advantageous to be closer to cover than your group members.

Second, individuals may become vigilant because they observe the same external stimulus from the environment (Pays et al. 2007a; Blanchard and Fritz 2007). This strategy explains synchronization from the standpoint of the predator, which would target stragglers (Beauchamp 2015). However, this hypothesis does not appear to explain synchronization of perch behavior in Barbary ground squirrels as we found that synchronization occurred when predators were and were not observed. Yet, we found an interaction between the proportion of time spent perched and predator presence with lower levels in synchronization when predators were present. This result may be driven by the lower sample size when predators were present.

Third, synchronization may also be explained by the behavior of the group. The squirrels perform perch behavior mainly in the morning and afternoon. So, it may be that it is the first behavior of all squirrels after emergence from their nest burrows in the morning, and that they become satiated within a couple of hours of foraging and then perch in the afternoon. Thus, the synchrony may be an emergent property of the ecology of the species.

Finally, we suggest another mechanism besides the copying effect and collective behavior, to explain synchronization of perch bouts: the presence of rock walls in the habitat. These rock walls prevent the squirrels from detecting terrestrial ambush predators when on the ground, as they cannot look on either side of the wall. Visual cues are essential for predator detection in diurnal species and, thus, for ground-dwelling sciurids, whereby predator detection is easier in an open-structured habitat (Ylönen and Brown 2007; Embar et al. 2011). Hence, to adequately detect ambush predators, i.e., feral cats, two Barbary ground squirrels may be necessary to look in both directions of the rock wall, which was supported by the result that in 40% of scans in absence of predators and in 60% of scans in presence of predators more than 1 squirrel was perched. Synchronization of perch bouts may also explain our earlier result that individual high-quality vigilance sometimes increased with group size (van der Marel et al. 2019). In the stingless bee, *Tetragonisca angustula*, guards are present to defend the nest entrance and these guards are evenly distributed on either side of the entrance to increase the view of their surroundings (Shackleton et al. 2018). Therefore, similar to synchronized vigilance in bees, perch behavior may simply be a form of extremely exclusive vigilance where individuals improve their level of detection by synchronizing vigilance bouts to watch ‘each other’s back’. Further research is needed to test whether perch behavior may function as a form of collective sentinel behavior and whether perched squirrels scan in opposite directions or different visual areas.

Some of our results were not as expected. Other studies have reported some persistent synchronization of vigilance, but the difference in proportions has often been small (Pays et al. 2007a, b, 2012; Öst and Tierala 2011; Podgórski et al. 2016; Wang et al. 2020), while in our study the difference was large. This larger difference in proportions may be explained by a group size effect on synchronization (Beauchamp 2015). Barbary ground squirrels appeared to show greater synchrony with increasing group size, which supports the copying effect. The group size effect can also be explained by dilution of predation risk (Pays et al. 2009), an individual’s body condition (Öst and Tierala 2011), and the amplification effect, where the group’s behavior amplifies an individual’s behavior (Krause and Ruxton 2002; Pays et al. 2009). Furthermore, the sample sizes that we used, observation period and sleeping burrow associations, may not be the optimal method to calculate coordination as Barbary ground squirrels show fission-fusion dynamics (van der Marel et al. 2020). Using observation period, we included all individuals in sight during a scan but this method may have resulted in pseudoreplication as individuals can be observed on multiple days. Using sleeping burrow associations, we prevented pseudoreplication but we excluded individuals that belonged to another sleeping burrow association, even though they were observed in the same scan, resulting in lower confidence of coordination. However, using both these extremes, we still obtained the same result, suggesting that we captured the real coordination pattern of Barbary ground squirrels.

## Conclusions

This study shows that quantitative testing of coordination is essential to describe a sentinel system. In addition, we demonstrated that multiple non-mutually exclusive hypotheses may explain synchronization of perch bouts in the invasive population of Barbary ground squirrels: Synchronized vigilance might occur as squirrels copy the behavior of group members (copying effect), as an artefact of the behavior of the group, and as an adaptation to anthropogenically altered habitats (watch each other’s back). Thus, synchronized vigilance may result in fitness benefits, particularly, in habitats where the field of view is obstructed by man-made structures.

## Acknowledgments

We acknowledge the landowners of our study sites, the IPNA-CSIC by providing a research vehicle, and the Cabildo of Fuerteventura for the use of the facilities and the help of its staff at the Estación Biológica de La Oliva. We thank the editors and 2 anonymous reviewers for their constructive feedback.

## Declarations

### Ethics approval

All applicable international, national, and institutional guidelines for the care and use of animals were followed. All procedures performed were in accordance with the ethical standards of the institution or practice at which the studies were conducted (University of Manitoba Animal Care and Use Committee, protocol no. F14-032, and the government of Fuerteventura, Cabildo Insular de Fuerteventura no. 14885).

### Availability of data and code

The datasets analyzed and code used during the current study are available from a Github repository (https://github.com/annemarievdmarel/coordinated-vigilance, van der Marel et al. 2021b).

### Authors’ contributions

All authors conceived the ideas and designed methodology; AM collected the data, analyzed the data, and led the writing of the manuscript; all authors contributed critically to the drafts and gave final approval for publication.

